# An Ensemble Learning Approach for Cancer Drug Prediction

**DOI:** 10.1101/2020.08.10.245142

**Authors:** Darsh Mandera, Anna Ritz

## Abstract

Predicting the response to a particular drug for specific cancer, despite known genetic mutations, still remains a huge challenge in modern oncology and precision medicine. Today, prescribing a drug for a cancer patient is based on a doctor’s analysis of various articles and previous clinical trials; it is an extremely time-consuming process. We developed a machine learning classifier to automatically predict a drug given a carcinogenic gene mutation profile. Using the Breast Invasive Carcinoma Dataset from The Cancer Genome Atlas (TCGA), the method first selects features from mutated genes and then applies K-Fold, Decision Tree, Random Forest and Ensemble Learning classifiers to predict best drugs. Ensemble Learning yielded prediction accuracy of 66% on the test set in predicting the correct drug. To validate that the model is general-purpose, Lung Adenocarcinoma (LUAD) data and Colorectal Adenocarcinoma (COADREAD) data from TCGA was trained and tested, yielding prediction accuracies 50% and 66% respectively. The resulting accuracy indicates a direct correlation between prediction accuracy and cancer data size. More importantly, the results of LUAD and COADREAD show that the implemented model is general purpose as it is able to achieve similar results across multiple cancer types. We further verified the validity of the model by implementing it on patients with unclear recovery status from the COADREAD dataset. In every case, the model predicted a drug that was administered to each patient. This method will offer oncologists significant time-saving compared to their current approach of extensive background research, and offers personalized patient care for cancer patients.

## Introduction

Almost one in six deaths occur due to cancer, and the number of new cancer cases is expected to rise by 70% in the next twenty years (“WHO | Cancer”, 2017). In today’s world, the process of prescribing a drug for a cancer patient is based on a doctor’s exploratory research through various scientific journals, such as the Cochrane Review, PubMed, et cetera, as well as previous clinical trials. This can become an extremely time-consuming and iterative process.

The principal goal of precision medicine is to treat patients with the treatments that are most likely to help the patients, based on their genome (“Precision Medicine in Cancer Treatment – National Cancer Institute”, 2017). In cancer, this entails predicting which treatment will be the most effective on a patient based on their genetic mutations. Machine learning, which is a branch of artificial intelligence that allows computers to “learn” from experience, has been used in various aspects of cancer prediction and prognosis (Simes, 1985; Maclin et al., 1991; Ciccheti, 1992; Petricoin and Liotta, 2004; Bocchi et al., 2004; Zhou et al., 2004; Dettling, 2004; McCarthy et al., 2004; Wang et al., 2005; Cruz and Wishart, 2007, Yasrebi et al., 2009, Yasrebi, 2016). Combined with a high volume and wide range of patients’ records consisting of gene data along with effectiveness of medications, machine learning will help predict the best way to administer the correct targeted medication for patients with cancer (Kourou et al., 2014; Bashiri et al., 2017). In today’s world, the majority of such established machine learning programs use data from cancer cell lines – sets of cancer cells that grow and divide in a laboratory. Still others use humanized mice models – human xenografts and immune cells grafted into lab mice (Kalamara, et el., 2018). Although these may be effective models, it is impossible for any model to fully capture the complexity of real patient data.

This project utilizes supervised machine learning and aims at creating a general-purpose machine learning classifier that can predict which drug will be effective for a cancer patient based on their genetic mutations, for any cancer type. Overall, this program uses patterns and correlations between a patient’s response to a drug and their genetic mutations in order to predict which drug will be effective on a given cancer patient, based on their genetic mutations.

## Methods

This project involved multiple steps including data wrangling, training the machine learning model, and testing the machine learning model (Figure 1).

**Figure 1:**
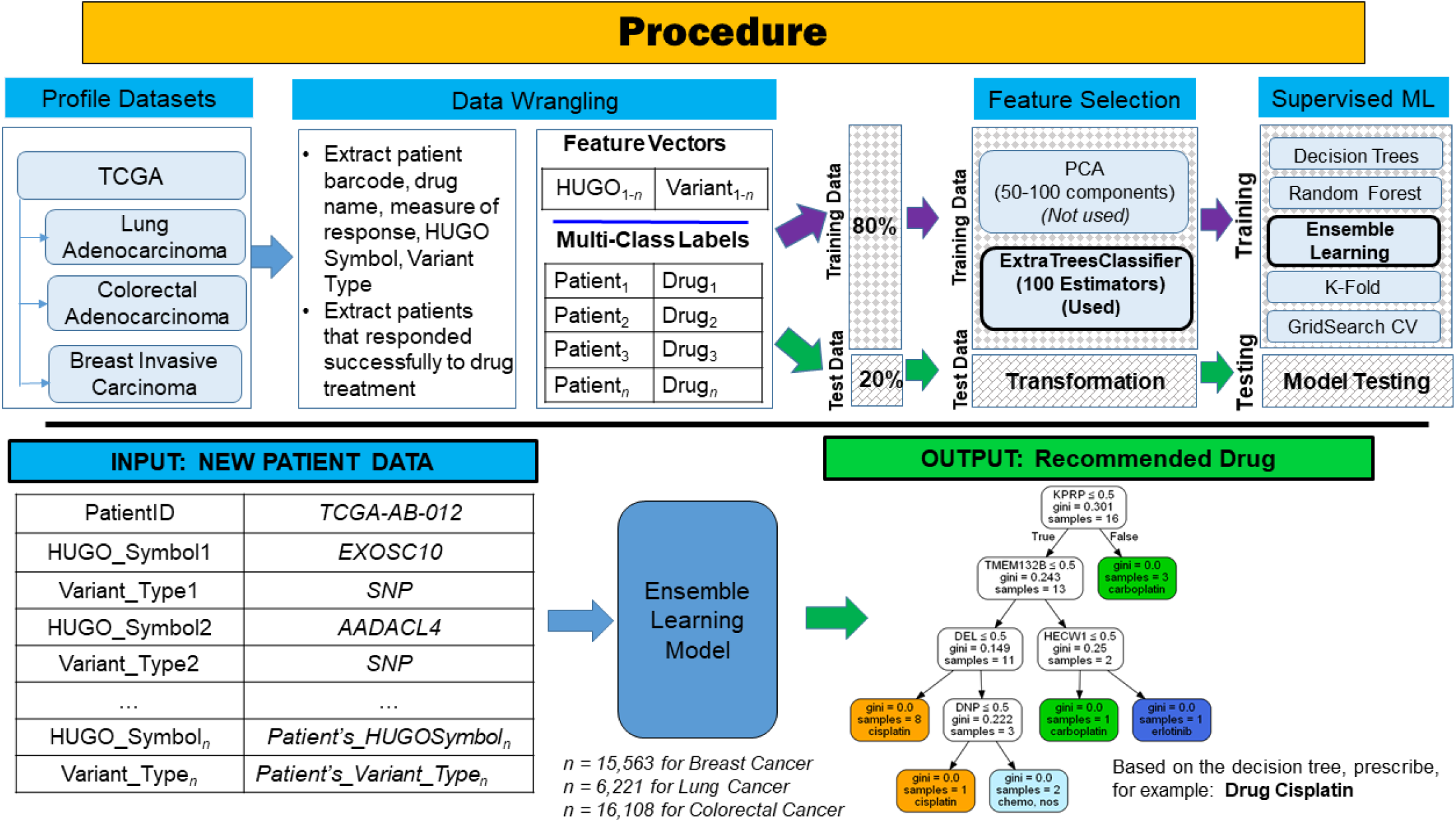
Various stages of the machine learning model training and prediction on various cancer datasets. Data from The Cancer Genome Atlas were formatted to extract features (mutations and variants) and multi-class drug labels for each patient. Then, we performed feature selection and supervised machine learning for four different classifiers. Once the model is trained, new patient data can be classified in the Ensemble Learning Model to recommend a drug for that patient.

The first stage of this project was data acquisition and data wrangling. The data was acquired from The Cancer Genome Atlas (TCGA), which is a collaboration between the National Cancer Institute (NCI) and National Human Genome Research Institute (NHGRI), and has generated comprehensive, multi-dimensional maps of the key genomic changes in 33 types of cancer (“The Cancer Genome Atlas Home Page,” n.d.). First, a Breast Invasive Carcinoma (BRCA) dataset with 982 patients was downloaded. Patient barcode (patient’s anonymous TCGA ID), drug names, measure of response (a patient’s response to a specific drug), HUGO Symbol (name of the gene where a genetic mutation occurs), and variant type (the type of genetic mutation that a patient has) were extracted from the BRCA dataset consisting of over 15,000 data types (features).

In machine learning, feature selection can be undertaken in order to remove features that do not contribute to the accuracy of the model. In this research, we implemented feature selection using the ExtraTreesClassifier on the BRCA dataset. The ExtraTreesClassifier implements an estimator that fits randomized decision trees on samples of the dataset in order to select features that have the highest impact on the final accuracy, allowing for reduction of overfitting. (“3.2.4.3.3. sklearn.ensemble.ExtraTreesClassifier”, 2019) Principal Component Analysis (PCA) was also tested for feature selection, but ExtraTreesClassifier resulted in a higher accuracy of 52%, while PCA returned an accuracy of 30%. The use of the ExtraTreesClassifier resulted in the identification of the top 20 features (various HUGO Symbols and Variant Types), and their impact on the accuracy of the program (Figure 2).

**Figure 2:**
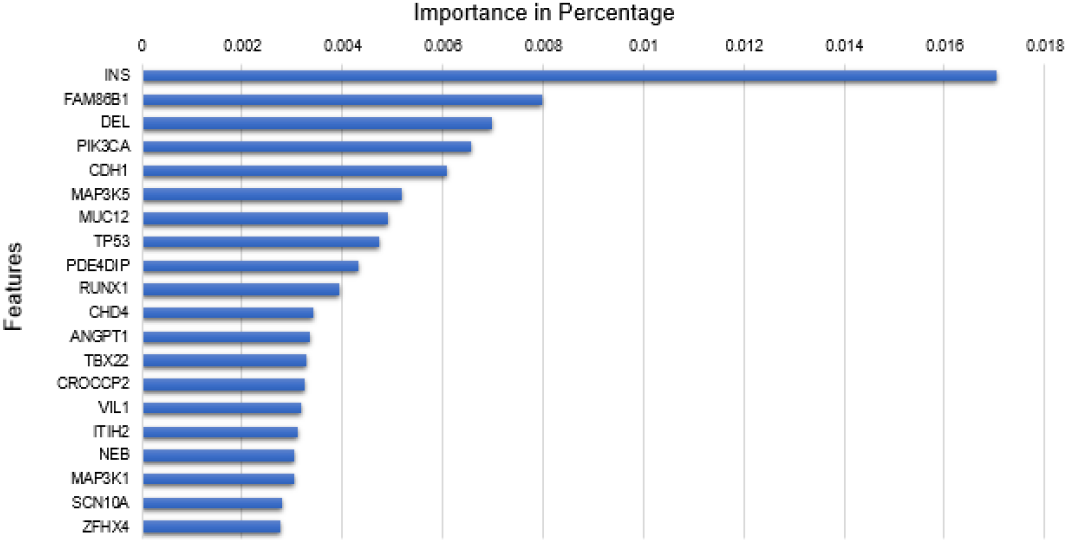
Top 20 features that were selected from the BRCA dataset.

The resulting BRCA dataset had 695 patients that had successfully responded to drugs (i.e. had a complete response) for their particular genetic mutation profile. For patients who were administered more than one drug, we chose the first drug that resulted in a complete response as the target. This dataset was then converted into two vectors: one containing all HUGO Symbols and Variant Types, and the other containing patient barcodes and drug names.

Once the data had been curated, the extracted dataset of 695 patients was split into a training section and a testing section using the train_test_split function in scikit-learn. 80% of the data was used for training, and the remaining 20% was used for testing, as an 80-20 split is a recommended best practice.

The training dataset was used to fit the model to learn which drug is the most effective for a set of mutated genes. On the training data with features, K-Fold and GridSearchCV were implemented for cross-validation. K-Fold, with a K-value selected to equal 5, split the training dataset into five equal folds. GridSearchCV was then implemented on each of these five folds as a hyperparameter optimization algorithm and selected the best parameters to use for prediction. The processes of train_test_split and cross-validation were repeated using four different classifiers, in order to identify which machine learning algorithms would result in the highest prediction accuracy. The following is a brief description of the four algorithms that we tested:

### DecisionTree Classifier

A decision tree outputs a tree-like structure that decides which drug a patient should receive based on their genetic mutations. The decision tree starts with a type of mutation and asks: *Does a patient have this mutation or not?* The tree then decides: *If not, then do not prescribe them any drug. If they do, prescribe Drug_X.* (“1.10 Decision Trees”, 2020)

### AdaBoost Classifier

The AdaBoostClassifier combines weak algorithms to form a strong prediction algorithm. AdaBoostClassifier works by training multiple base estimators on the training data, and then using a weighted voter among those estimators in order to select a final classification. In this study, we chose our base estimator to be RandomForest. (“sklearn.ensemble.AdaBoostClassifier”, 2019)

### One vs. Rest Classifier

This classifier takes a unique approach on classifying an item. The program selects one drug, then combines the rest of the drugs into a single class. Then, it selects a certain combination of HUGO Symbols and Variant Types and decides whether the combination belongs in the one selected class or in the large combined class. Then, the algorithm selects a new drug to isolate. The algorithm repeats this process until all possible permutations are executed. (“sklearn.multiclass.OneVsRestClassifier”, 2020)

### K-Nearest-Neighbor Classifier

This last algorithm works by learning from *k* closest neighbors to an unclassified patient. It then chooses a classification for the given patient based on the class of its *k* closest neighbors. (“1.6. Nearest Neighbors”, 2019).

In order to validate that the final machine learning model was general-purpose, a Lung Adenocarcinoma (LUAD) dataset with 178 patients and 33 drugs and a Colorectal Adenocarcinoma (COADREAD) dataset with 223 patients and 23 drugs were downloaded from TCGA, and these datasets underwent a similar data wrangling process as the BRCA dataset. LUAD and COADREAD data was then inputted into the machine learning model. The model was designed and implemented in Python 3.6 using scikit-learn 0.19.1, pandas 0.20.3, and pydot 1.2.3 in order to predict the best drug for all cancer types. All code and data are available at https://github.com/annaritz/cancer-drug-pred.

## Results

Overall, AdaBoostClassifier and OneVsRestClassifier yielded two of the highest training prediction accuracies of 85% and 83% respectively, and they were implemented in the testing phase. The BRCA test data of 139 patients was scored on the trained model based on AdaBoostClassifier and OneVsRestClassifier. AdaBoostClassifier yielded an accuracy of 63%, while OneVsRestClassifier yielded the highest testing accuracy of 66%. Therefore, OneVsRestClassifier was used in the final model.

The final machine learning model was tested on BRCA data as well as LUAD and COADREAD datasets to evaluate it as a general-purpose program. On the BRCA data, the program was able to predict effective drugs on a patient with a training accuracy of 83% and a testing accuracy of 66%. On the LUAD dataset, the program achieved a training accuracy of 55% and a testing accuracy of 50%. On the COADREAD dataset, the training accuracy and testing accuracy were both 66%. The reason that the accuracy values vary highly is due to the fact that some datasets (e.g. LUAD) may be sparse or have a smaller sample size compared to other datasets. (Figure 3).

**Figure 3:**
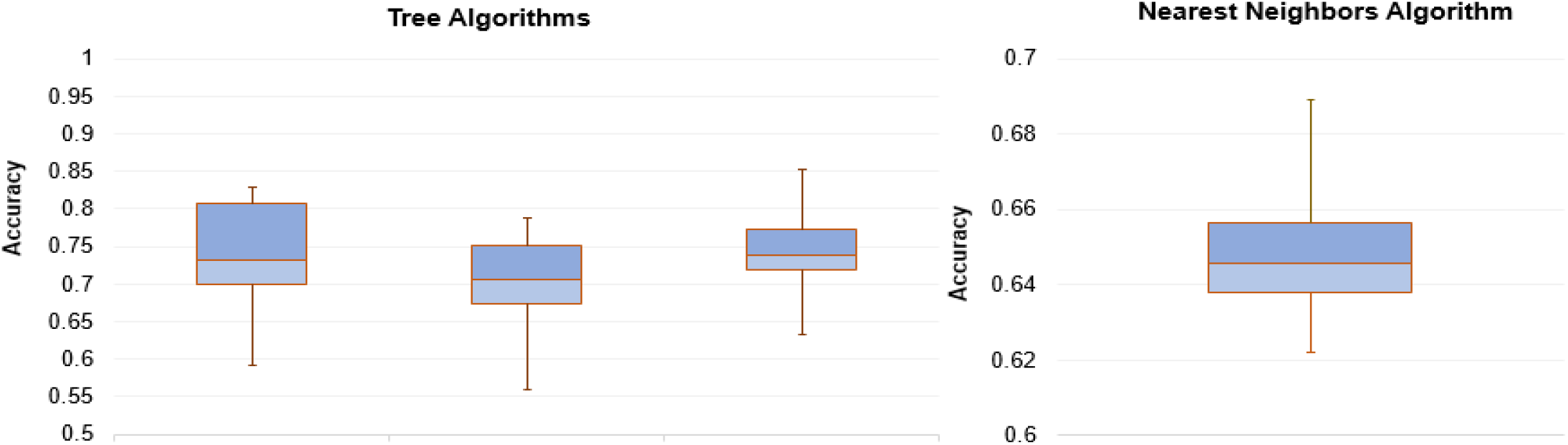
The variance of accuracy of the program tested on the BRCA dataset.

In this study, the most commonly selected drugs for Breast Invasive Carcinoma were cyclophosphamide (sometimes labeled as Cytoxan), tamoxifen, doxorubicin (labeled as Adriamycin), and anastrozole (labeled as Arimidex). The DecisionTree Classifier for BRCA is shown in Figure 4. Cyclophosphamide is often used to treat all types of breast cancer, in addition to other cancers like ovarian cancer and leukemia. As an alkylating agent, cyclophosphamide slows the growth of breast cancer cells or, in some cases, completely halts the growth. (“Cyclophosphamide: MedicinePlus Drug Information”, 2018; Cocquyt et al., 2003). Tamoxifen, another commonly selected drug for BRCA, works by binding to estrogen receptors in order to prevent estrogen binding (“Hormone Therapy for Breast Cancer Fact Sheet - National Cancer Institute”, 2017). It too has been used to treat breast cancer (Baum et al., 2002; Houghton et al., 2003; Silva et al., 2014). Doxorubicin and anastrozole both work by slowing down cancer cell growth; the former by blocking topo isomorase 2 and the latter by blocking estrogen receptors (“Doxorubicin (Adryiamycin) | Cancer Drugs | Cancer Research UK”, 2017; “Anastrozole”, 2018). Previous research has demonstrated these drugs’ efficacy in BRCA patients (Baselga et al., 1998; Baum et al., 2002; Chae et al, 2016).

**Figure 4:**
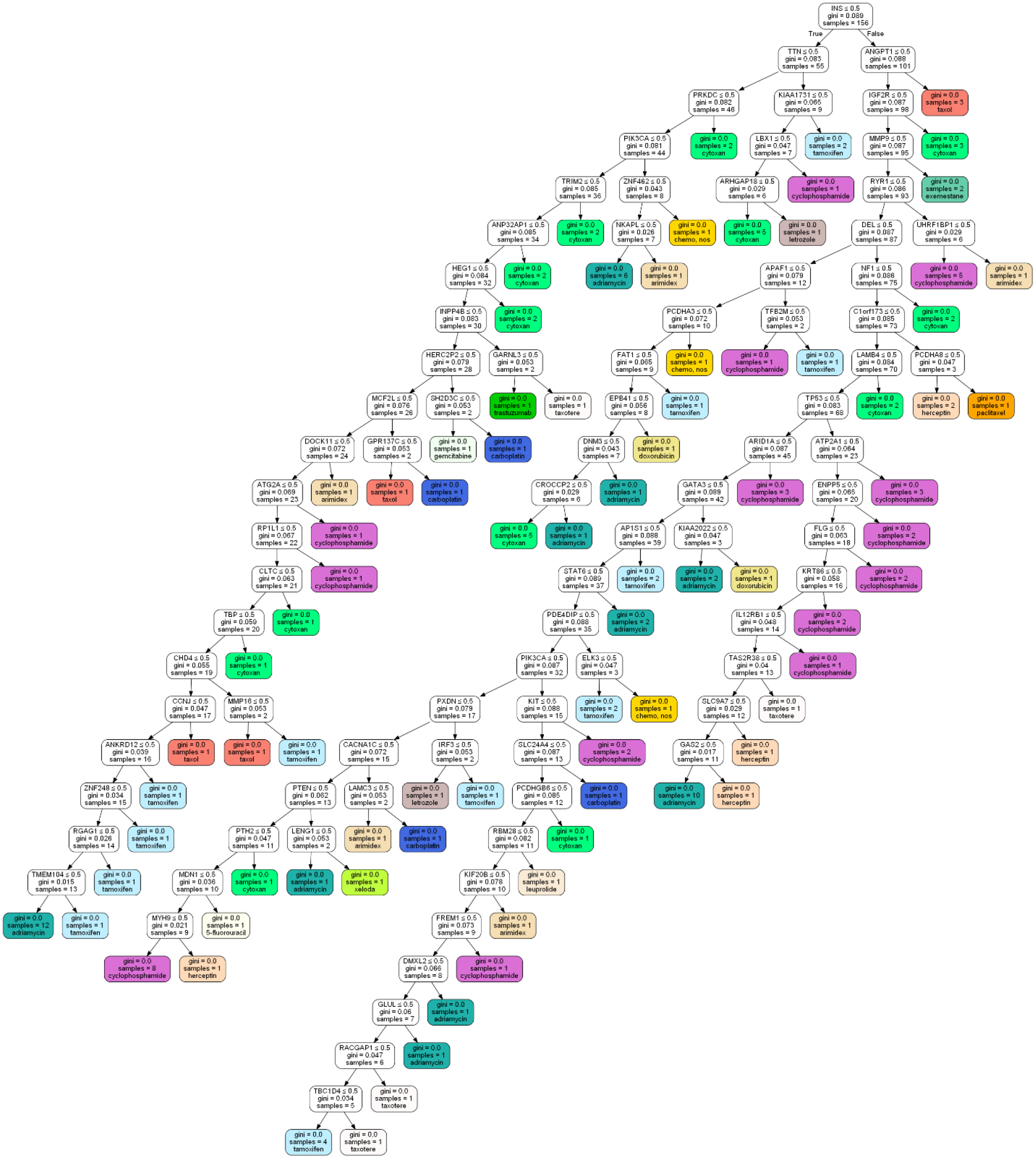
A decision tree generated based on the BRCA dataset that determines which drug a patient should receive based on their genetic mutations.

In order to further test the validity of the classifier, 200 patients from the COADREAD dataset who did not recover from their disease were selected. These patients varied in age and sex. Fifty-two percent of these unclassified patients were male, while the remaining 48% were female. Two percent were under 18, 2% were between the ages of 19 and 40, 15% were between the ages of 41 and 64, and 81% were over 65. For each of these patients, the classifier predicted a drug that had been administered by the clinic but did not help the patient recover. For example, for the patient TCGA-A6-2677, the classifier predicted to prescribe the drug oxaliplatin, which had been administered by the clinic. The most commonly selected drugs for unclassified patients were oxaliplatin, 5-fluorouracil, and bevacizumab, which are among the most commonly prescribed drugs for colorectal adenocarcinoma (“Targeted Therapy Drugs for Colorectal Cancer”, 2018; “Chemotherapy for Colorectal Cancer”, 2018). Overall, these results show that the classifier is able to predict drugs that are commonly prescribed by oncologists in clinical settings.

## Discussion

Predicting which drugs will be most effective on a cancer patient based on their genetic mutations is an important goal of precision medicine. A general-purpose machine learning classifier was designed and implemented to predict targeted drug for cancer based on patients’ genetic mutations. Overall, the program that was created can be used by doctors to identify which drugs to prescribe to a cancer patient for targeted therapy. While the model is able to predict drugs similarly to a doctor’s judgement, it is important to note that the current tool still cannot distinguish between when drugs will be effective and when patients will not respond.

The next steps to this project will be to further validate the general purpose of the machine learning model by testing it against other cancer types (e.g. Head and Neck, Prostrate, et cetera), ideally with larger datasets. Also, expanding the machine learning model to use more dimensions such as patient lifestyle and location of the mutated gene to narrow the scope and predict with higher accuracy (90% or more) would be ideal. Another plan could be to create a new machine learning model that predicts partial or complete efficacy of more than one drug, that is, model the program as a multi-class problem.

## Author Contributions

D.M. began this project in 2017, and sought the mentorship of A.R. D.M. and A.R. formulated the prediction problem, and D.M. carried out all experiments. D.M. wrote the initial drafts of the manuscript, and D.M. and A.R. worked on the manuscript together. D.M. also presented this research at the Reed College Poster Session as well as the Oregon Health and Science University Research Week in 2018, and the Rocky Mountains Bioinformatics Conference in 2019.

